# Combining genetic crosses and pool targeted DNA-seq for untangling genomic variations associated with resistance to multiple insecticides in the dengue vector *Aedes aegypti*

**DOI:** 10.1101/575753

**Authors:** Julien Cattel, Frédéric Faucon, Bastien Lepéron, Stéphanie Sherpa, Marie Monchal, Lucie Grillet, Thierry Gaude, Frederic Laporte, Isabelle Dusfour, Stéphane Reynaud, Jean-Philippe David

**Affiliations:** Laboratoire d’Ecologie Alpine (LECA), UMR 5553 CNRS - Université Grenoble-Alpes, Grenoble, France; Institut Pasteur de la Guyane, Cayenne, France

**Keywords:** Insecticide resistance, Mosquito, *Aedes aegypti*, Copy Number Variations, Polymorphism, Complex phenotype, Detoxification enzymes, Cytochromes P450.

## Abstract

In addition to combating vector-borne diseases, studying the adaptation of mosquitoes to insecticides provides a remarkable example of evolution-in-action driving the selection of complex phenotypes. Indeed, most resistant mosquito populations show multi-resistance phenotypes as a consequence of the variety of insecticides employed and of the complexity of selected resistance mechanisms. Such complexity makes challenging the identification of alleles conferring resistance to specific insecticides and prevents the development of molecular assays to track them in the field. Here we showed that combining simple genetic crosses with pool targeted DNA-seq can enhance the specificity of resistance allele’s detection while maintaining experimental work and sequencing effort at reasonable levels. A multi-resistant population of the mosquito *Aedes aegypti* was exposed to three distinct insecticides (deltamethrin, bendiocarb and fenitrothion) and survivors to each insecticide were crossed with a susceptible strain to generate 3 distinct lines. F2 individuals from each line were then segregated with 2 insecticide doses. Bioassays supported the improved segregation of resistance alleles between lines. Hundreds of genes covering all detoxifying enzymes and insecticide targets together with more than 7,000 intergenic regions equally spread over mosquito genome were sequenced from pools of F0 and F2 individuals unexposed or surviving insecticide. Differential coverage analysis identified 39 detoxification enzymes showing an increased gene copy number in association with resistance. Combining an allele frequency filtering approach with a Bayesian F_ST_-based genome scan identified multiple genomic regions showing strong selection signatures together with 50 non-synonymous variations associated with resistance. This study provides a simple and cost-effective approach to improve the segregation of resistant alleles in multi-resistant populations while reducing false positives frequently arising when comparing populations showing divergent genetic backgrounds. The identification of these insecticide resistance markers paves the way for the design of novel DNA-based resistance tracking assays.

**Author summary:** In addition to combating vector-borne diseases, understanding how mosquitoes adapt to insecticides provides a remarkable example of evolution-in-action. However, the variety of insecticides used and the complexity of adaptive mechanisms make it difficult to identify the genetic changes conferring resistance to each insecticide. Here we combined simple controlled crosses with mass DNA sequencing for enhancing the specificity of resistance gene detection. A multi-resistant mosquito population was exposed to three distinct insecticides and survivors were crossed with a susceptible strain to generate 3 distinct mosquito lines. Individuals from the second generation of each line were then segregated based on their resistance to each insecticide. Bioassays supported the improved segregation of genetic resistance markers between lines. Hundreds of genes potentially involved in resistance together with thousands non-genic regions equally spread over mosquito genome were sequenced from individuals from each line. Genomic analyses identified detoxification enzymes showing an increased gene copy number in association with resistance and multiple genomic regions showing strong selection signatures and carrying point mutations associated with resistance. Such approach improves the specificity of resistance gene detection in field mosquito populations resisting to multiple insecticides and paves the way for the design of novel DNA-based resistance tracking tools.

## Introduction

Natural populations experience a variety of selective pressures, leading to the accumulation of locally adaptive features and the expression of complex phenotypes [1]. Environmental changes driven by man-made disturbances can alter the course of selection, by inducing novel, particularly strong and sometimes unpredictable selective pressures. Understanding how natural populations respond to rapid environmental changes has become a major goal, and an increasing number of studies reported adaptive changes on very short timescales [2-4]. Resistance of insects to insecticides is a key example of rapid evolution under novel and strong selective pressures associated with human activities. This adaptive phenotype has evolved quickly and independently in a large number of taxa [5]. However, natural resistant populations often exhibit complex resistance phenotypes as a consequence of the variety of insecticides used, the intensity of the selection pressures, and the selection of mechanisms conferring resistance to multiple insecticides, making the identification of resistance alleles challenging [6, 7]. Besides contributing to the understanding of rapid adaptation and the origins of complex traits, deciphering the complexity of insecticide resistance mechanisms could help improving risk assessments and management strategies [8].

Among taxa of serious economic and medical importance, mosquitoes are vectors of numerous human viruses and pathogens representing a major threat for public health worldwide [9]. Among them, *Aedes aegypti* is of particular importance because of its wide distribution [10] and its capacity to transmit several major arboviral diseases including Yellow Fever, Dengue, Zika fever and Chikungunya fever. Although efforts are invested in developing novel vaccines and strategies to prevent arbovirus transmission, the use of chemical insecticides remains the cornerstone of arboviral diseases control. However, as for malaria vectors, decades of chemical treatments have led to the selection and spread of resistance in this mosquito species. Insecticide resistance is now widespread in *Ae. aegypti* and affects all insecticide families used in public health [11], often leading to reduced vector control efficacy [12-14]. Although attempts are made to develop alternative arbovirus control strategies [15] their large scale implementation will require decades. Until this, characterizing molecular mechanisms underlying resistance is crucial for tracking down resistance alleles and improving resistance management strategies [16].

Resistance of mosquitoes to chemical insecticides can be the consequence of various mechanisms, such as non-synonymous mutations affecting the protein targeted by insecticides, a lower insecticide penetration, its sequestration, or its biodegradation often called metabolic resistance [6, 17]. In *Ae. aegypti*, resistance to pyrethroids, the main insecticide family used against mosquitoes, is mainly the consequence of target-site mutations affecting the voltage-gated sodium channel targeted by these insecticides (Knock Down Resistance ‘*kdr’* mutations) and metabolic mechanisms [11, 18]. Several *kdr* mutations have been identified in this species and the causal association between the V410L, S989P, V1016G/I and F1534C mutations and pyrethroid resistance has been confirmed [19-23]. Most of these mutations can be genotyped on individual mosquitoes by PCR-based assays, providing essential allele frequency data for resistance management. Conversely, metabolic resistance is far less understood in *Ae. aegypti* although this type of resistance is frequent and often accounts for a significant part of the resistance phenotype [6]. Such resistance mechanism is caused by an increased activity of detoxification enzymes. These detoxification enzymes include cytochrome P450 monooxygenases (P450s or *CYPs* for genes), carboxy/cholinesterases (CCEs), glutathione S-transferases (GSTs) and UDP- glycosyl-transferases (UDPGTs) although other families can be involved [17, 18, 24]. Their high diversity (∼ 300 genes in *Ae. aegypti*) and the complexity of biodegradation pathways make challenging the identification of those conferring resistance to a specific insecticide. Theoretically, metabolic resistance can be the consequence of an increased expression of one or multiple detoxification enzymes metabolizing the insecticide and/or the selection of variants showing a higher insecticide metabolism rate due to conformal modifications. As over expression is frequently associated with over transcription, most candidate genes were identified based on their differential transcription in resistant populations as compared to susceptible counterparts using transcriptomics [11, 18, 24, 25]. Although these approaches identified several detoxification enzymes involved in insecticide biodegradation, they mostly failed to pinpoint the underlying genomic changes, thus impairing the high-throughput tracking of metabolic resistance alleles in natural populations. The application of powerful genomic tools has improved the understanding of the genetic bases of metabolic resistance in *Ae. aegypti*. Using deep targeted DNA sequencing (targeted DNA-seq) Faucon et al. [26] identified genomic variations associated with resistance to the pyrethroid deltamethrin in multiple populations sampled from different continents. This study identified several detoxification enzymes affected by Copy Number Variations (CNV) and non-synonymous variations in association with insecticide resistance. Cross-comparing these genomic data with transcriptomic data obtained from RNA-seq confirmed the central role of CNV in the over-expression of detoxification enzymes associated with resistance in this species [27]. However, this study used natural resistant populations displaying multi-resistance phenotypes, thus not allowing to properly discriminating between alleles specifically associated with resistance to the insecticide in question and those associated with resistance to other insecticides. Furthermore, this approach did not allow breaking up the genetic linkages between the genomic variations identified, thus potentially leading to false positives.

In this context, the present study aimed at better understanding the origin of complex insecticide resistance phenotypes in the mosquito *Ae. aegypti*. More precisely, we combined genetic crosses and targeted DNA-seq in an attempt to identify genomic variations specifically associated with resistance to distinct insecticides in a multi-resistant *Ae. aegypti* population. After exposure to three insecticides of distinct chemical families (the pyrethroid deltamethrin, the organophosphate fenitrothion and the carbamate bendiocarb), survivors to each insecticide were crossed with a susceptible strain to generate three F2 lines. Each F2 line was then phenotyped with two increasing doses of its respective insecticide and survivors were used to identify CNV and polymorphism variations associated with resistance in hundreds of target genes including all detoxification enzymes and insecticide target proteins. In addition, the capture of thousands of intergenic regions regularly distributed over mosquito genome also allowed crossing up these data with a genome-wide screening of selection signatures associated with resistance. Overall, this study contributes to improve our understanding of the complex genomic bases of metabolic resistance to insecticides and paves the way for the design of novel insecticide resistance tracking tools in this major arbovirus vector.

## Results

### Insecticide resistance levels

The population used in this study consisted in a composite population representative of multiple natural *Ae. aegypti* populations collected from French Guiana (see methods). Bioassays performed on the initial composite population (F0 Guy-R) confirmed its high resistance to the pyrethroid insecticide deltamethrin with resistance ratio (RR_50_) over 316-fold as compared to the susceptible strain Bora-Bora (Fig. 1A). This population also showed moderate resistance to the carbamate bendiocarb and the organophosphate fenitrothion with RR_50_ of 14-fold and 3-fold respectively. As expected, resistance to each insecticide decreased after crossing F0 survivors to each insecticide with the susceptible strain with F1 resistance ratios decreasing to 25-fold, 7-fold and 2-fold for deltamethrin, bendiocarb and fenitrothion respectively. Deltamethrin resistance was even lower in F2 (10-fold) while fenitrothion resistance remains low (1.8-fold) and bendiocarb resistance slightly increased to 10-fold. Assessing the cross resistance of each line to all insecticides confirmed the partial segregation of resistance alleles after controlled crosses (Fig. 1B). Although F2 individuals from each line showed a higher survival when exposed to its respective insecticide, this trend was only significant for deltamethrin in link with the lower resistance to other insecticides.

**Fig 1.**
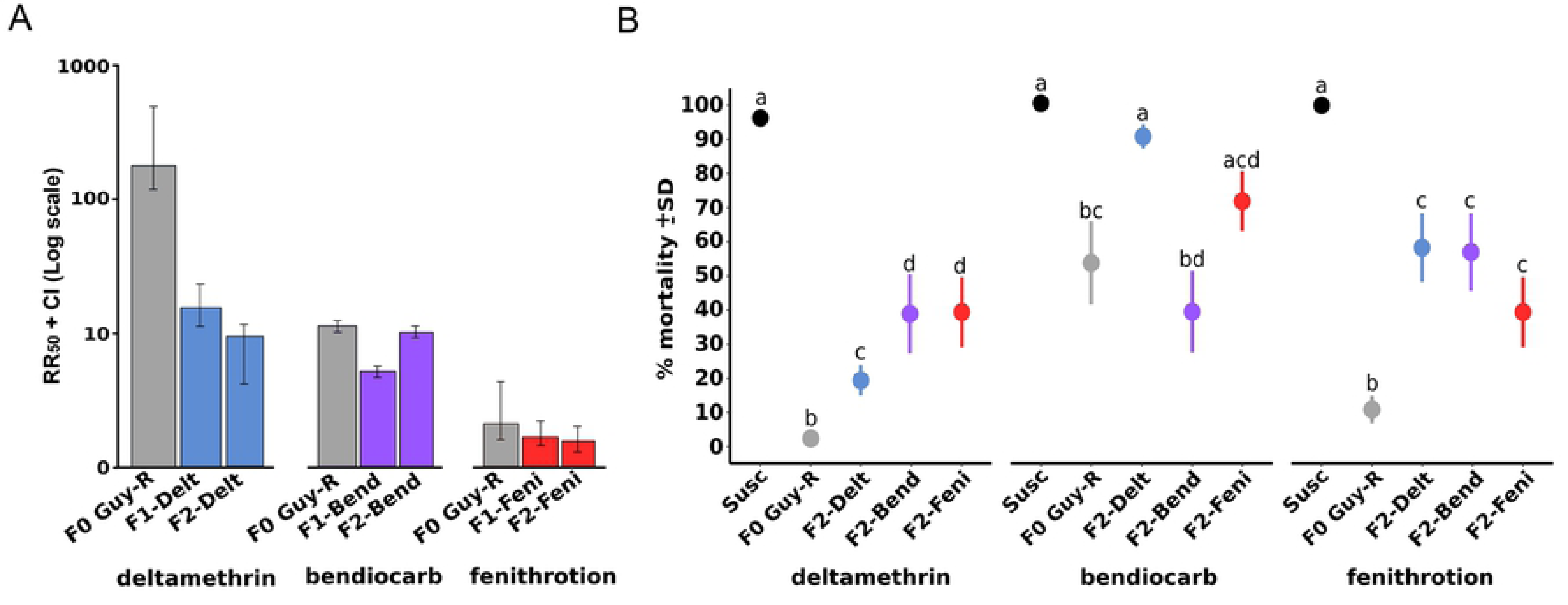
Insecticide resistance Levels. Resistance levels of the different lines to the three insecticides deltamethrin, bendiocarb and fenitrothion. Black: susceptible strain, Grey: F0 Guy-R composite population, Blue: Delt line, Purple: Bend line, Red: Feni line. A: Resistance levels of each line to its respective insecticide at the F0 Guy-R, F1 and F2 generations. Resistance levels are expressed as LD_50_ ± 95% CI. B: Cross-resistance profiles of each line to all insecticides at the F2 generation. Cross-resistance levels are expressed as % mortality ± SD to a single insecticide dose. For each insecticide, letters indicate statistical similarity or dissimilarity between lines (GLM family = binomial, N≥4).

### Target-site resistance mutations

Assessing kdr mutations frequencies from targeted DNA-seq reads data confirmed the high frequency of the three kdr mutations V410L, V1016I and F1534C in F0 Guy-R composite population corroborating its high deltamethrin resistance level (Fig. 2). Exposing F0 Guy-R individuals to the LD_80_ of each insecticide did not segregate these mutations in survivors. The segregation of kdr mutations became more evident in F2 individuals with higher allele frequencies observed in F2 individuals of the Delt line surviving to high dose of deltamethrin. As expected because genetically constrained by two successive mutation events in *Ae. aegypti* [28], the acetylcholinesterase G119S mutation conferring resistance to organophosphates and carbamates in other species was not detected for DNA-seq reads data. Validation of kdr mutation frequencies on individual mosquitoes by qPCR confirmed the robustness of allele frequencies estimated from DNA-seq although moderate discrepancies were observed when the number of genotyped mosquitoes was low (S1 Fig.).

**Fig 2.**
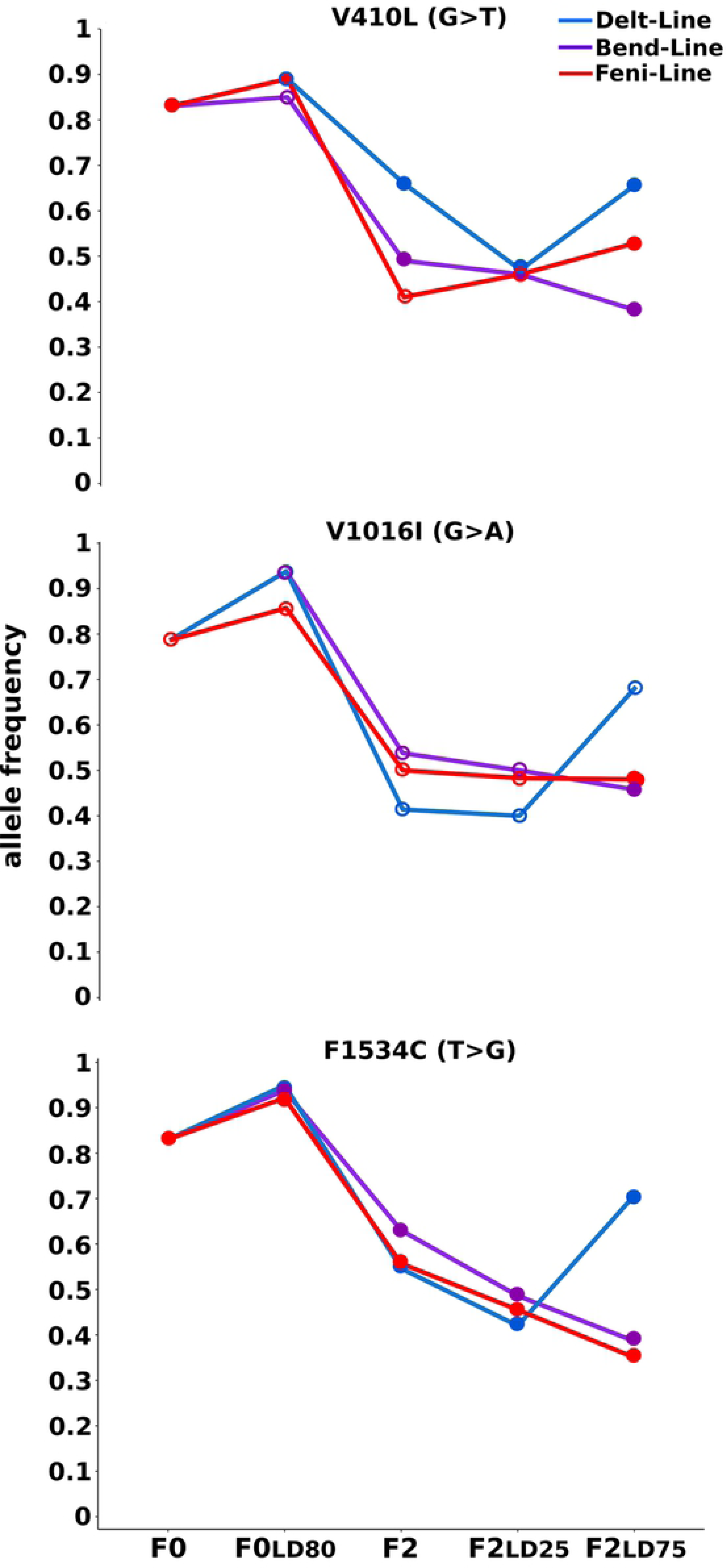
Evolution of kdr mutations frequencies. Allelic frequencies variations of the three kdr mutations V410L, V1016I and F1534C initially present in the F0 Guy-R population in each line. Allele frequencies were inferred from the number of sequencing reads supporting each allele at each locus. Empty dots indicate conditions with read coverage < 30.

### Gene copy number variations

Over 49.5 % of sequenced reads were successfully mapped to AaegL5 exome allowing the detection of 1,317 exonic regions (719 distinct genes) showing a minimum length of 45 bp and a coverage between 30 and 800 reads/bp in all conditions (median = 94.1 reads/bp). Filtering genes based on their expected CNV profiles across F0 and F2 conditions in each line (see methods for filtering conditions) allowed identifying 39 detoxification genes affected by CNV in association with insecticide resistance (Fig. 3 and S1 Table). Although the resistance level of the Delt line was high, more CNV were detected in the two other lines likely due to the contribution of kdr mutations in the deltamethrin resistance phenotype. This trend was also observed for CNV intensity as most genes identified in the Delt line showed a lower CNV increase in F2 individuals surviving high dose of insecticide as compared to those identified in the Bend and Feni lines.

**Fig 3.**
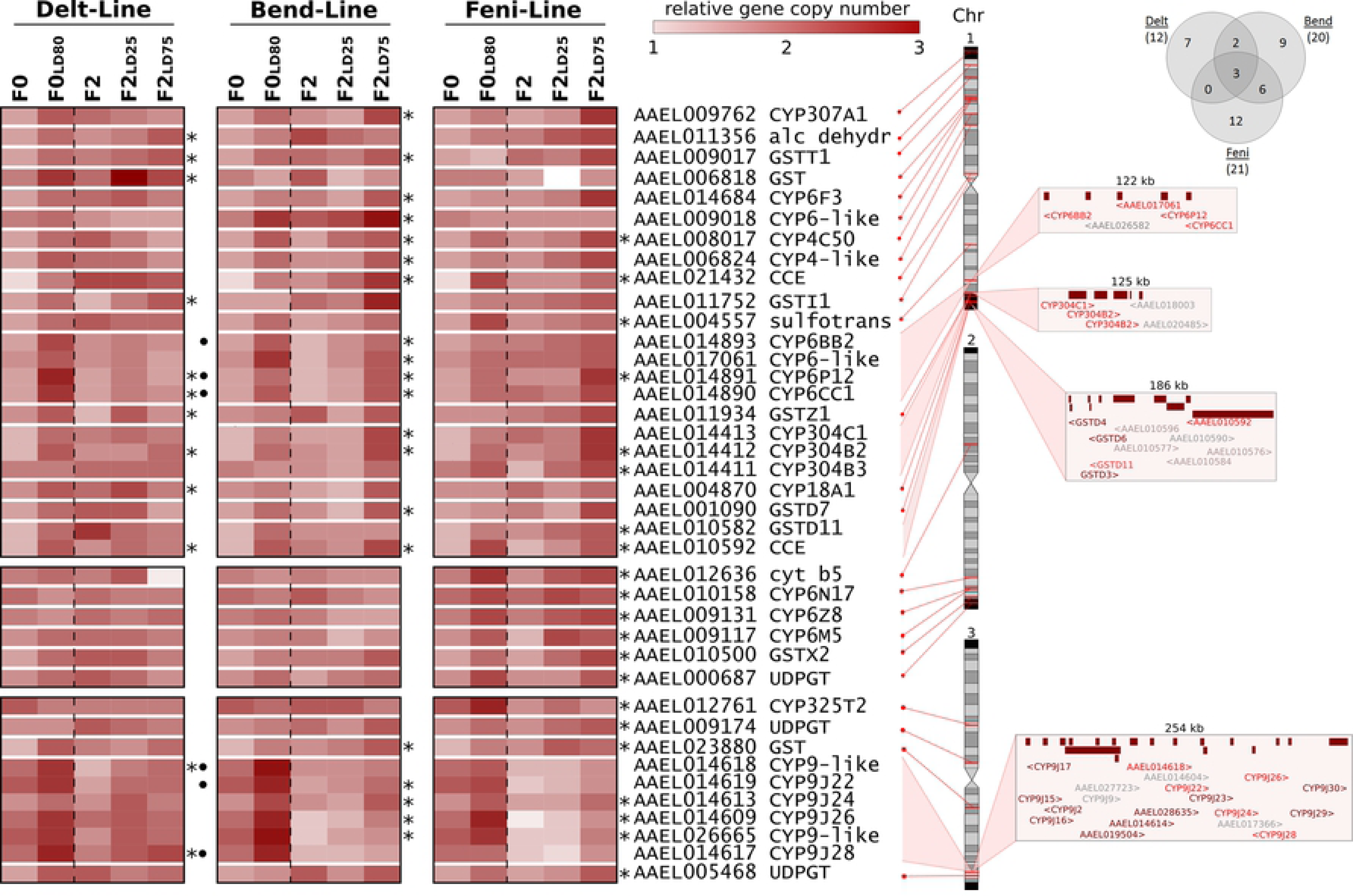
Gene copy number variations associated with insecticide resistance. For each line, genes affected by CNV associated with resistance are indicated by stars. CNV data obtained from F0 Guy-R are repeated for each line for better clarity. Black dots indicate genes previously identified as affected by CNV associated with deltamethrin resistance in Faucon et al. [26]. The genomic location of each gene on chromosomes and gene clusters architecture is shown on the right (red: CNV associated with resistance, brown: CNV not associated with resistance, grey: genes not included in the targeted regions). The Venn diagram indicates the number of genes affected by CNV associated with resistance in each line.

Among genes affected by CNV, 11 were associated with resistance to multiple insecticides including the P450s *CYP6P12* and *CYP304B2* and the CCE *AAEL010592* being associated with resistance to all insecticides. Genes affected by CNV in association with deltamethrin resistance included 6 P450s, 4 GSTs, 1 alcohol dehydrogenase and 1 CCE. Among them, 2 CYP6 genes belonging to a cluster of P450s in chromosome 1 and 2 CYP9Js belonging to a large cluster of P450s on chromosome 3 were previously identified as affected by CNV associated with deltamethrin resistance [26]. Genes affected by CNV in association with bendiocarb resistance included 15 P450s, 3 GSTs and 2 CCEs. P450s included all genes of the CYP6 cluster located on chromosome 1 and 4 genes of the large CYP9J cluster located on chromosome 3 but also several genes from the CYP4, CYP6 and CYP304 families. The CYP6-like *AAEL009018* located on chromosome 1 was specifically associated with bendiocarb resistance with a marked CNV increase in both F0 and F2 survivors. Genes affected by CNV in association with fenitrothion resistance included 11 P450s, 3 GSTs, 3 glycosyl-transferases (UDPGTs), 2 CCEs and 1 sulfotransferase. Genes specifically associated with fenitrothion resistance included the sulfotransferase *AAEL004557* and the P450 *CYP304B3* on chromosome 1 and the UDPGT *AAEL005468* located at the end of chromosome 3, but most genes were located at the end of chromosome 2 (*CYP6N17*, *CYP6Z8*, *CYP6M5*, *GSTX2*, UDPGT *AAEL000687*).

### Selection imprints and polymorphisms

Over 85% of sequenced reads were successfully mapped to AaegL5 genome allowing the detection of more than 40,000 variations. Among them, 24,714 were polymorphic across conditions and passed quality and coverage filters (S2 Table). These variations were mostly substitutions (96.6%) and were mostly located in targeted regions (70.4 %). The mean distance between two variations was ∼50 kb. Filtering these polymorphisms based on their expected frequency variations across F0 and F2 conditions in each line (see methods for filtering conditions) allowed identifying 302 (1.23%) differential polymorphisms associated with insecticide resistance across all lines. Most of them were line-specific with only three of them shared between the Feni and Bend lines. Combining allele frequency filtering with a F_ST_-based Bayesian approach allowed detecting multiple genomic regions carrying differential polymorphisms and showing low Q values in both F0 and F2 samples in any line (Fig. 4 and S3 Table). Most of them were located in proximity of genes potentially involved in metabolic resistance and included genes carrying non-synonymous variations associated with resistance (see below). Among these regions, 5 were located on chromosome 1, including two GST and 1 P450 clusters. The P450 cluster (3 CYP304 genes at ∼287 Mb) showed a pronounced selection signature for the Delt and Bend lines while the GST cluster located at ∼300 Mb appeared more associated with the Bend line. Several regions were also detected on chromosome 2. One ABC transporter cluster (4 genes at ∼90 Mb), 1 sulfotransferase cluster (2 genes at 134.15 Mb) and 2 CCE clusters (6 genes at ∼174 Mb and 4 genes at ∼214 Mb) showed strong selection signature in the Feni line. The large GST cluster (15 GSTE genes at ∼351.5 Mb) was associated with both Feni and Delt lines while the large CYP6 cluster (16 genes at ∼419.2 Mb) showed a strong selection signature in all lines. Among regions identified in chromosome 3, the 2 large P450 clusters (21 CYP325 genes at ∼111.6 Mb and 18 CYP9J genes at ∼368.5 Mb) and the sulfotransferase cluster (6 genes at 396.8 Mb) were detected in all lines. Despite the 81 polymorphisms detected in the voltage-gated sodium channel gene carrying kdr mutations (gene *AAEL023266* ∼316 Mb), only a moderate selection signature was detected in this region mainly in the Delt and Bend lines.

**Fig 4.**
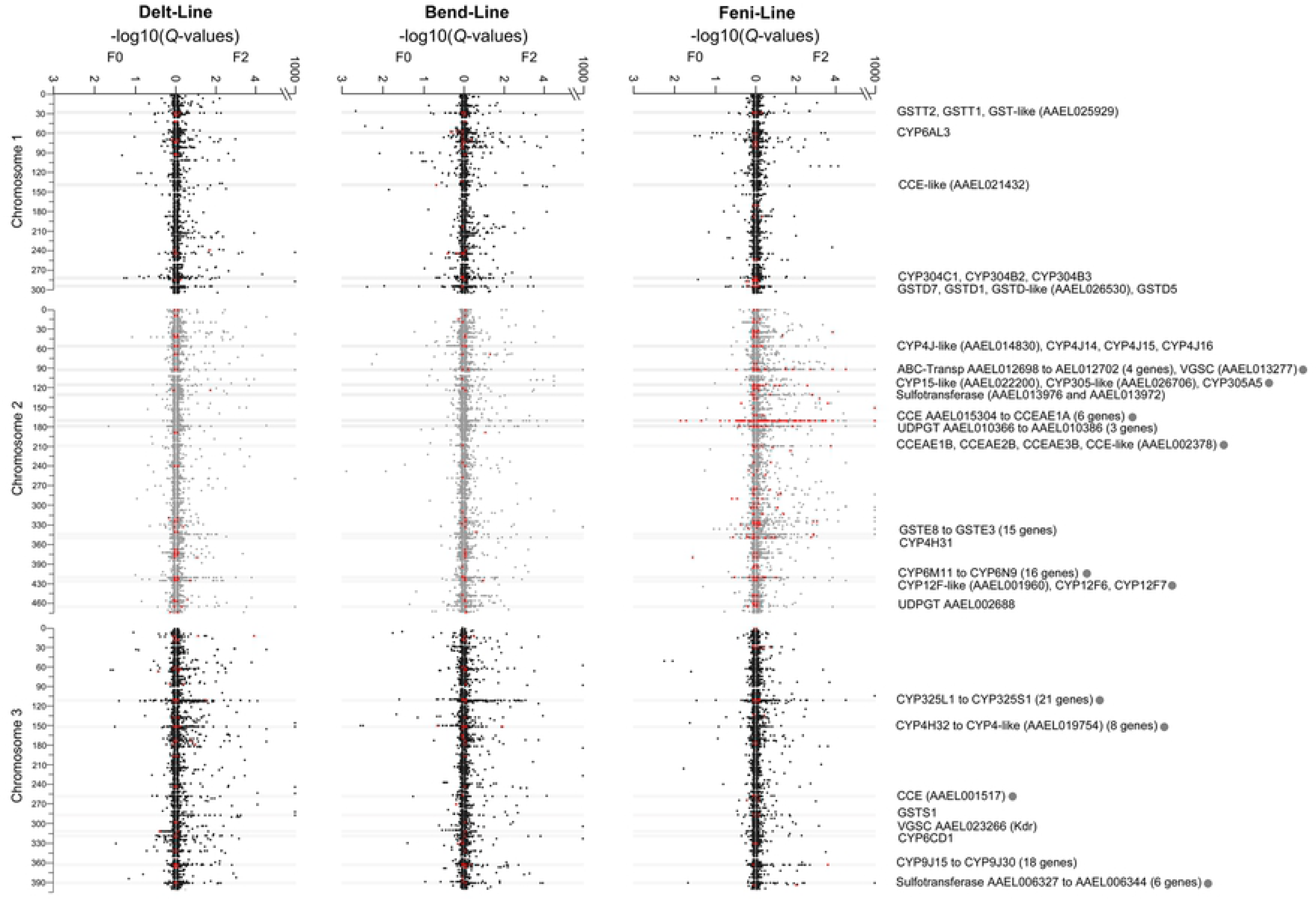
Selection signatures associated with insecticide resistance. For each line, region under selection were identified based on the presence of differential polymorphisms from allele frequency filtering (red dots, see methods for filtering conditions) and the presence of loci displaying low Q values in both F0 and F2 conditions. Q values were computed separately in F0 (left arm) and F2 conditions (right arm) and Q values = 0 were fixed at 10^−1000^ for better clarity. Horizontal grey lines indicate genomic regions showing strong selection signatures associated with resistance. For each region the names of potential resistance genes located within a 50 kb range are indicated. Grey dots indicate genomic regions carrying genes affected by differential non-synonymous variations associated with resistance as shown in Fig 5.

Among all differential polymorphisms associated with resistance identified, 50 were non-synonymous (Fig. 5 and S3 Table). Most of them affected detoxification genes located in genomic regions showing strong selection signatures. All of them were line-specific except the I324V mutation identified in the alcohol dehydrogenase gene *AAEL026142* found associated with resistance in both Bend and Feni lines. Seven were associated with resistance in the Delt line. These affected the alcohol dehydrogenase *AAEL020054* and the P450 *AAEL001960* in chromosome 2 together with 4 clustered P450s from the CYP325 family and 1 sulfotransferase in chromosome 3. Ten were associated with resistance in the Bend line affecting 8 distinct genes: 5 on chromosome 1, affecting the alcohol dehydrogenase *AAEL026142*, the P450s *CYP9AE1* and *CYP329B1*, and *GSTD6*; 2 on chromosome 2, affecting 2 CYP6 belonging to a large cluster of P450s (*CYP6M9* and *CYP6N13*); and 3 on chromosome 3, affecting the P450s *CYP4K3* (2 variations) and *CYP6AG3*. Finally, more than 30 non-synonymous variations were associated with resistance in the Feni line affecting 21 distinct genes. On chromosome 1, this included the alcohol dehydrogenase *AAEL026142* and *GSTD1* for which a coding frameshift was negatively associated with resistance. Multiple isolated genes were affected on chromosome 2, including 1 ABC transporter, a few P450s and 1 UDPGT. Two CCE clusters located within regions showing strong selection signatures (at ∼174 Mb and ∼214 Mb) were also affected with the first cluster being affected by a total of 17 point mutations. Three P450s (*CYP6M11*, *CYP6Y3* and *CYP6N13*) located within a large CYP6 cluster (at ∼419 Mb) were also affected on chromosome 2. Finally, only the P450 *CYP325T1* and the CCE (*AAEL001517*) were affected on chromosome 3.

**Fig 5.**
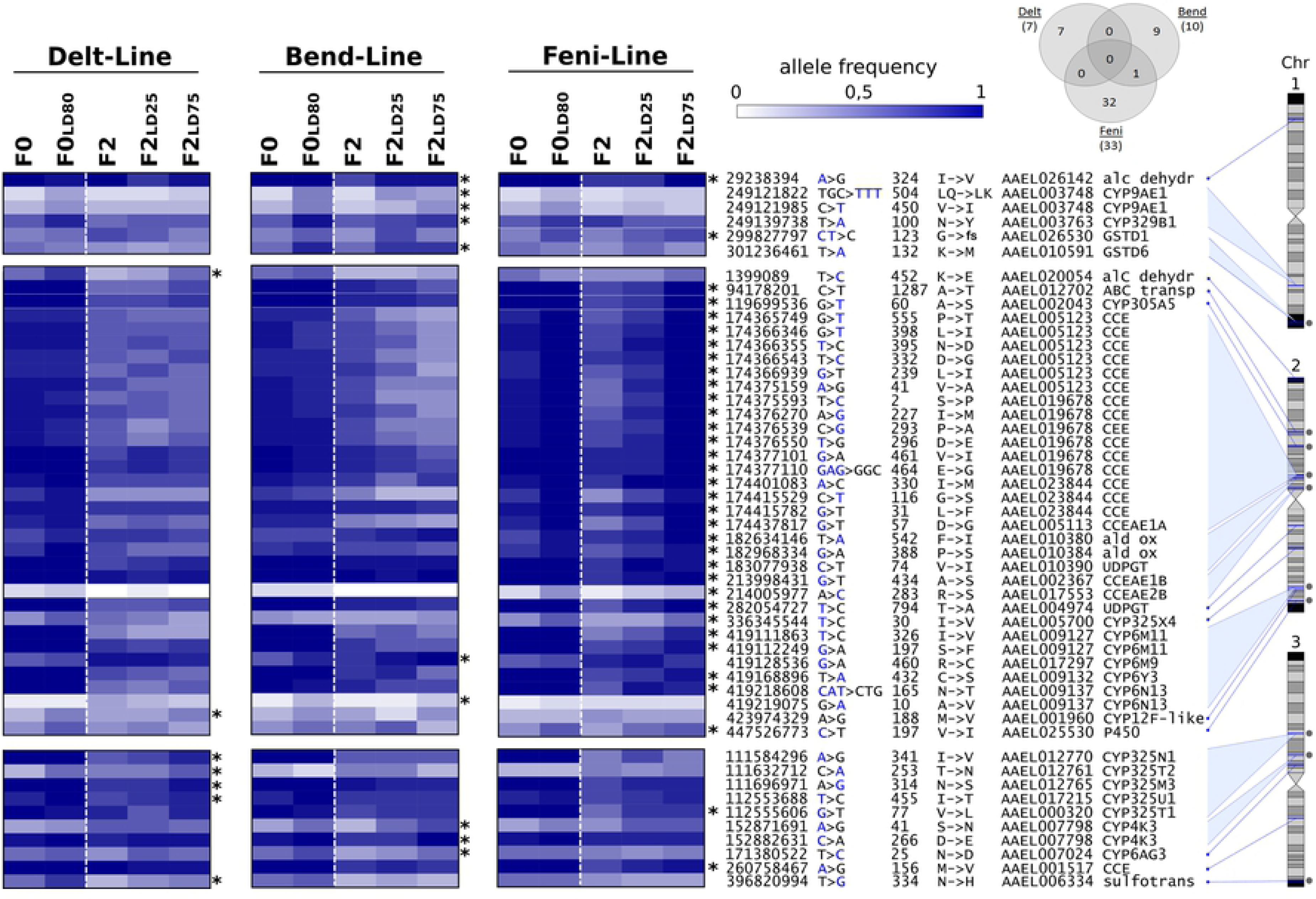
Non-synonymous polymorphisms associated with insecticide resistance. The Venn diagram indicates the number of non-synonymous polymorphisms identified in each line. The heat map shows allele frequencies in each condition. For each variation the allele represented on the heat map is indicated in blue. Allele frequency data obtained from F0 Guy-R are repeated for each line for better clarity. For each line, stars indicate non-synonymous polymorphisms associated with resistance based on their expected allele frequency profile across all conditions. Grey dots indicate non synonymous polymorphisms located in genomic regions showing strong selection signatures as shown in Fig 4. For each variation the following annotations are shown: genomic location on chromosome, reference nucleotide > variant nucleotide, amino acid position, amino acid change (fs= frame shift), gene accession number, gene name.

## Discussion

Natural populations experience a variety of selective pressures often leading to the expression of complex adaptive phenotypes. In mosquitoes transmitting human diseases, an over-reliance on chemical control has resulted in the rapid selection and spread of alleles conferring resistance to various insecticides, often leading to multi-resistance phenotypes [6, 11, 29]. As opposed to target-site mutations which are specific to a given insecticide mode of action, the complexity and redundancy of insect detoxification systems underlying metabolic resistance make it less predictable and can lead to the selection of various and multiple resistance alleles depending on the local context [6, 30]. Most insecticide resistance studies using field mosquito populations focused on resistance mechanisms to a given insecticide or a given chemical family. However, these kinds of studies did not fully discriminate alleles associated with resistance to different insecticides, which may lead to false positives. In this context, the present study attempted at demonstrating that the identification of resistance alleles can be improved by combining simple genetic crosses and targeted DNA pool sequencing, while maintaining experimental work and sequencing costs at reasonable levels.

### Using controlled crosses for segregating resistance alleles to different insecticides

Bioassays confirmed the high resistance of the F0 Guy-R composite population from French Guiana to deltamethrin and its moderate resistance to the two other insecticides. As expected, the introgression of susceptible alleles by controlled crosses strongly reduced deltamethrin resistance in F2 individuals. This suggests that resistance alleles approaching fixation in the initial F0 population were less present in F2 individuals, thus facilitating their dose-response segregation. This was confirmed by the strong decrease of kdr mutations frequencies observed between F0 and F2 individuals in each line. Cross-resistance patterns obtained from F2 individuals supported the partial segregation of resistant alleles between each line. This segregation was also supported by the divergent kdr mutations frequency patterns observed from F0 to F2 conditions between lines. Nevertheless, an incomplete segregation of resistance alleles was expected as only two generations of recombination are likely not enough to break out genetic associations between alleles conferring resistance to distinct insecticides in all individuals. Such partial segregation may also indicate that particular genes/alleles are contributing to resistance to multiple insecticides. This was previously shown in *Anopheles* mosquitoes and *Drosophila melanogaster* where particular detoxification enzymes have been shown to metabolize multiple insecticides from different chemical families [31-34].

### Detoxification enzymes CNV associated with resistance

Metabolic resistance is frequently associated with the over-expression of detoxification enzymes having the ability to degrade and/or sequester insecticides [6, 17]. Although changes in gene expression can be caused by cis- or trans-mediated transcriptional or post-transcriptional regulation, CNV may also impact gene expression. Initially associated with organophosphate resistance in *Culex pipiens* [35], recent genomic studies confirmed the involvement of CNV in metabolic resistance to various insecticides in mosquitoes [26, 27, 36]. Such key role of CNV in metabolic resistance is not surprising as the locus mutation rate is far higher for CNV than for mutation [37]. In addition CNV events are favored by the presence of transposable elements which account for a large part of most mosquito genomes [∼50% in Ae. aegypti genome, 38]. Furthermore, CNV have a direct impact on gene expression level (*i.e.* gene dosage effect) without necessarily altering protein function [39], suggesting that advantageous duplications have moderate costs and can be rapidly selected in natural populations undertaking strong insecticide pressures. Finally, it has been shown in yeast that a specific environmental change can stimulate the occurrence of CNV affecting genes involved in this specific adaptation [40]. Considering that the proposed transcriptional mechanism depends on promoter activity and that detoxification enzymes are frequently inducible by insecticides, such mechanisms may have also favored the selection of CNV-mediated metabolic resistance to insecticides in mosquitoes.

Our study identified 39 detoxification genes affected by CNV in association with resistance to insecticides. Although the F0 Guy-R composite population from French Guiana exhibits a high resistance to the pyrethroid deltamethrin, only few CNV were found associated with resistance to this insecticide and most of them did not show a strong dose-response in F2 individuals. Such low CNV signal was likely caused by the presence of kdr mutations which are known to be of significant importance in deltamethrin resistance in *Ae. aegypti* [11, 18, 22]. This was confirmed by the high kdr mutations frequencies found in individuals surviving to high dose of deltamethrin. However, our data also supported the added value of an increased gene copy number of detoxification enzymes in deltamethrin resistance, especially in F0 survivors for which kdr mutations are nearly fixed. This trend was particularly apparent for CNV affecting a cluster of 4 CYP6 genes located on chromosome 1 and for a large cluster of CYP9J genes located on chromosome 3. Indeed the over-expression of P450s belonging to these two gene clusters was previously associated with pyrethroid resistance in *Ae. aegypti* [11, 18, 24] and in *Aedes albopictus* [41]. Some of these genes, such as *CYP6BB2*, *CYP9J28* and *CYP9J32* have been functionally validated as able to metabolize pyrethroid insecticides [42, 43] and CNV contributing to the over-expression of these P450s were previously identified in deltamethrin-resistant populations [26, 27]. Finally, the role of the over-expression of other detoxification enzymes such as GSTs in pyrethroid resistance has been previously suggested [44-47]. These findings are in accordance with our identification of CNV affecting the microsomal GST *AAEL006818* and *GSTI1* in association with deltamethrin resistance.

As compared to deltamethrin, more genes were affected by CNV associated with bendiocarb and fenitrothion resistance. Indeed, even though resistance levels to these two insecticides were lower, the absence of resistance mutations affecting the target of these insecticides in *Ae. aegypti* because of genetic constraints [28] may have strengthened the association of CNV with resistance.

Among genes affected by CNV associated with bendiocarb resistance, the CYP6 *AAEL009018* located on chromosome 1 showed a strong and specific dose-response association with bendiocarb survival in both F0 and F2. The over-transcription of this gene was previously identified in multi-resistant populations from the Caribbean [48] but also in Malaysian populations showing cross-resistance between pyrethroids and carbamates [49]. The weak association of this gene with deltamethrin and fenitrothion resistance observed in our study supports its role in carbamate resistance. Similarly, CNV affecting the CYP6 *AAEL017061* showed a strong and specific association with bendiocarb resistance. The role of this gene in resistance is supported by the strong association of CNV affecting its *An. gambiae* orthologue *CYP6AA1* with insecticide resistance [50] but also by the capacity of *An. funestus CYP6AA1* to metabolize bendiocarb [51].

Several CNV affecting various detoxification enzymes were associated with resistance to fenitrothion. The genes *CYP6N17*, *CYP6Z8* and *CYP6M5*, *GSTX2* and the UDPGT *AAEL000687* were specifically associated with fenitrothion resistance. Noteworthy, orthologous P450s, GSTs and UDPGTs were also found highly over-transcribed in a Greek *Aedes albopictus* strain selected with the organophosphate temephos [52] supporting their potential contribution to organophosphate resistance. The amplification of a CCE cluster known to play a key role in temephos resistance [53] was not detected in our study, most likely because this CCE amplification is not distributed in French Guiana as confirmed by previous CNV data [26, 27].

Overall, the present study supports the contribution of CNV in the over-expression of detoxification enzymes conferring insecticide resistance in mosquitoes. Although deciphering their genomic architecture and their spatial dynamics in natural populations would require further work, these CNV represent promising DNA markers for designing novel high-throughput molecular assays to track metabolic resistance in the field.

### Selection signatures and non-synonymous variations associated with resistance

The combination of allele frequency filtering and F_ST_-based selection signature detection allowed identifying several genomic regions associated with insecticide resistance. These regions are often located in close proximity to detoxification genes involved in insecticide metabolism or previously found over-expressed in resistant populations [11, 18], supporting the robustness of our approach. However, a few regions showing strong selection signatures were identified near genes rarely associated with resistance in *Ae. aegypti*. This included multiple P450s from the CYP325, CYP4 and CYP12 families but also GSTs, UDPGTs, ABC-transporters and sulfotransferases, which may all be potentially involved in insecticide metabolism pathways. Most of these regions are located far from known resistance loci and included detoxification genes carrying non-synonymous variations associated with resistance. This suggests they may represent additional resistance loci possibly linked to the selection of particular detoxification enzyme variants. Indeed, even though most studies on metabolic resistance focused on the identification of over-expressed detoxification genes, the selection of particular variants leading to an increased insecticide metabolism rate can also contribute to the overall resistance phenotype as demonstrated in the malaria vector *An. funestus* [31, 54]. Among genes carrying non-synonymous variations associated with resistance, the deletion leading to a frame-shift coding in *GSTD1* is of particular interest. Indeed, as expected if this GST contributes to insecticide metabolism, the functional allele was specifically associated with resistance to fenithrothion in both F0 and F1 conditions. This enzyme has been shown to catalyze DDT dechloration and to be expressed in detoxification tissues in *An. gambiae* [55, 56], supporting its role in insecticide resistance. Also of interest are the multiple non-synonymous variations associated with fenitrothion resistance affecting a cluster of CCE genes located at 174 Mb in chromosome 2, of which one (*CCEae3A*, *AAEL023844*) has been shown to sequester and metabolize the organophosphate temephos in both *Ae. aegypti* and *Ae. albopictus* [53]. However, previous studies associated temephos resistance to the over-expression of this CCE gene through increased gene copy number [26, 57] while no CNV was detected for this gene in the present study. This suggests that the selection of carboxylesterase variants may also contribute to organophosphate resistance. This hypothesis is also supported by the previous identification of point mutations in *CCEae3A* for which docking simulations predicted an impact on temephos binding [57]. Although none of these mutations were found associated with resistance in our study, some mutations identified in these CCEs are located near the catalytic triad (*e.g.* I330M in *CCEae3A* and D332G in *AEL005123*) or the active site (*e.g.* P293A in *AAEL019678*). Altogether, the present study allowed identifying multiple detoxification genes located in genomic regions under selection and carrying non-synonymous mutations associated with resistance. Although further work is required to validate their association with the phenotype, the present study paves the way for better understanding their **c**ontribution in insecticide resistance.

## Conclusions

Although insecticide resistance has often been described as a monogenic adaptation in response to a strong selection pressure, it frequently results from the accumulation of multiple physiological and metabolic changes often leading to complex phenotypes. Because of their nature, target-site mutations are usually well characterized in mosquitoes and can typically be genotyped by simple PCR-based molecular assays [11, 18]. In contrast, genomic changes associated with metabolic resistance are far more difficult to characterize for various reasons: First, metabolic resistance alleles frequently co-occur with target-site mutations, thus weakening their association with the overall resistance phenotype. Second, the complexity and redundancy of insect detoxification pathways can lead to the selection of multiple and diverse alleles through local adaptation. Third, increased insecticide metabolism can be the consequence of multiple and additive genetic changes including non-synonymous polymorphisms causing structural changes of detoxification enzymes but also up-regulation and increased gene copy number, both enhancing their expression. Although massive parallel sequencing appears as a powerful tool for untangling the complexity of the genetic bases of metabolic resistance, it association with a well-thought experimental design can help reducing both false negatives and false positives. Here, we demonstrated that combining simple genetic crosses with pool targeted DNA-seq can enhance resistance allele segregation and produce high coverage sequence data for identifying metabolic resistance alleles while maintaining experimental work and costs at an acceptable level. Our results also suggest that eliminating the effect of target-site mutations by controlled crosses or gene editing should improve the power of genotype-phenotype association studies targeting metabolic resistance alleles. Considering the global threat of insecticide resistance on vector control and the decades that will be necessary for the full deployment of insecticide-free strategies, untangling the genetic bases of insecticide resistance still represents a challenge for controlling vector-borne diseases.

## Materials and methods

### Ethics statement

Blood feeding of adult mosquitoes was performed on mice. Mice were maintained in the animal house of the federative structure Environmental and Systems Biology (BEeSy) of Grenoble-Alpes University agreed by the French Ministry of animal welfare (agreement n° B 38 421 10 001) and used in accordance to European Union laws (directive 2010/63/UE). The use of animals for this study was approved by the ethic committee ComEth Grenoble-C2EA-12 mandated by the French Ministry of higher Education and Research (MENESR).

### Mosquitoes

The multi-resistant composite *Ae. aegypti* population from French Guiana used in this study consisted in a pool of 6 natural populations collected in 2016 in the following localities: Cayenne (North-East), Sinnamary (North-East), Saint-Laurent du Maroni (North), Apatou (North-West), Maripasoula (West) and Saint-Georges (East). Each population was collected as larvae from up to 5 breeding sites located within a 5 km range. These populations were separately raised to the adult stage and blood fed to generate adults of the next generation. The composite population was then created by pooling 1000 virgin adults of both sexes from each population and breeding them together for 3 generations without insecticide selection in order to homogenize genetic backgrounds. The resulting composite population (F0 Guy-R) was used for controlled crosses.

### Controlled crosses

Batches of virgin F0 Guy-R females were exposed to a dose killing 80% of individuals (LD_80_) of 3 insecticides belonging to distinct chemical families: the pyrethroid deltamethrin, the organophosphate fenitrothion and the carbamate bendiocarb (same exposure conditions as for bioassays, see below). Females surviving to each insecticide were then crossed with the fully susceptible strain Bora-Bora (Susc) in order to create three distinct lines (Fig. 6). For each line, controlled crosses were repeated twice and consisted in mass-crossing 100 virgin females having survived insecticide exposure (F0-Delt_LD80_, F0-Bend_LD80_, F0-Feni_LD80_) with an equal number of virgin males from the susceptible strain. For each line, F1 individuals were allowed to reproduce freely and blood fed in order to generate F2 individuals. F2 individuals from each line were then segregated based on their resistance phenotype by exposing 3 days-old females to two increasing doses of their respective insecticide (LD_25_ and LD_75_). F0 and F2 individuals from each line, unexposed and surviving to insecticides, were used for molecular analyses (Fig. 6).

**Fig 6.**
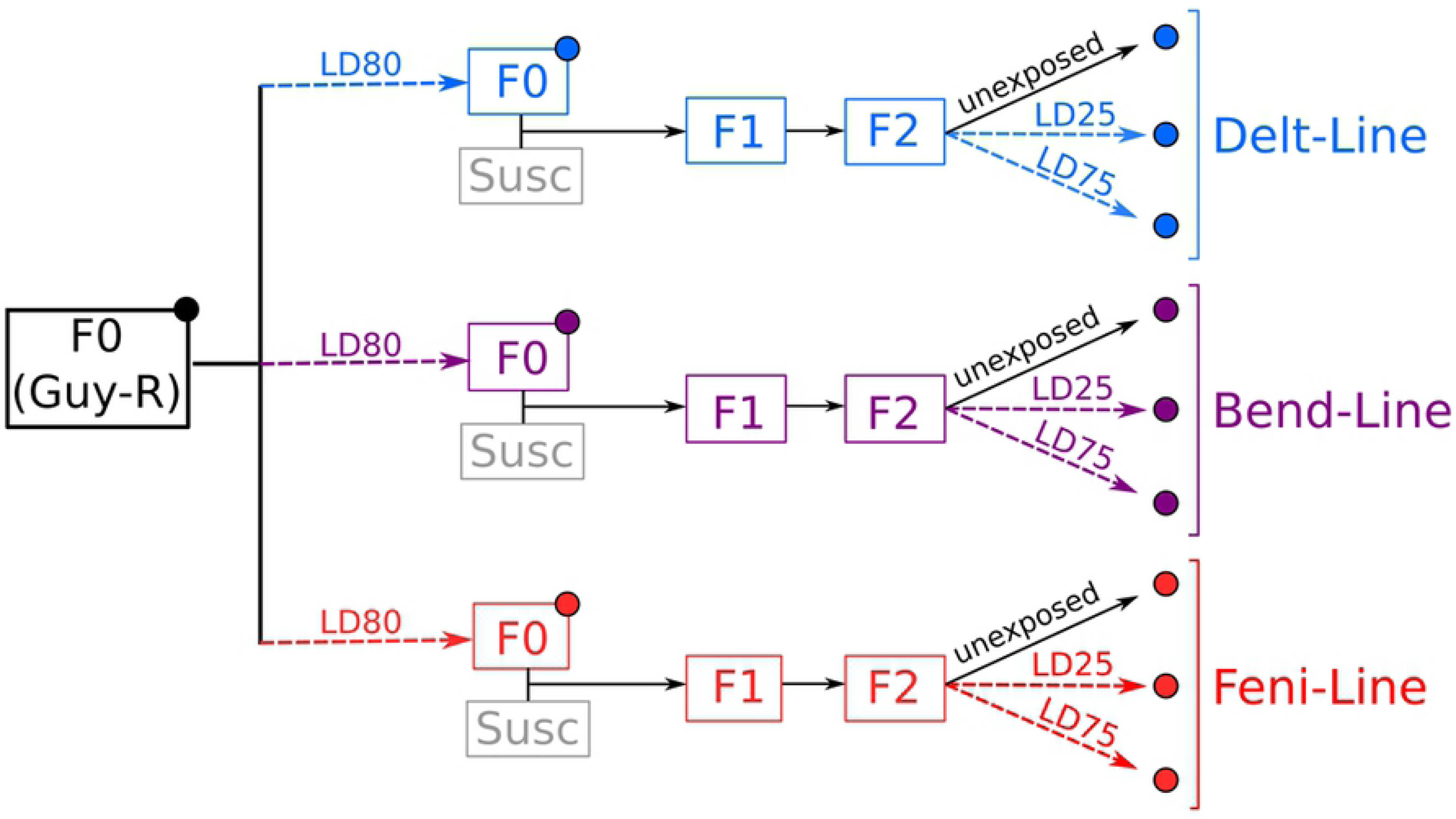
Experimental design overview. Insecticide survival segregation steps are shown as dashed arrows with the corresponding lethal dose (LD) indicated. Colors indicate insecticide lines (blue: Delt-line, purple: Bend-line, red: Feni-line). The initial resistant composite population (F0 Guy-R) and the susceptible strain (Susc) are shown in black and grey respectively. Large dots indicate samples used for targeted DNA-seq.

### Bioassays

All bioassays were performed on 3 days-old non blood fed females using WHO tests tubes equipped with insecticide-impregnated filter papers following WHO guidelines [58]. First, dose-response bioassays with deltamethrin, fenithrothion and bendiocarb were performed on the F0 Guy-R composite population to assess its resistance phenotype and identify the LD_80_ of each insecticide to be used for the initial F_0_ segregation. These bioassays were conducted with at least five doses of deltamethrin (0.05 % to 1%), fenitrothion (0.0125% to 0.4%) and bendioacarb (0.2 to 2%) and an exposure time of 60 min. At least five batches of 20 mosquitoes were used for each insecticide dose. Mortality data were recorded after a 24h recovery time and submitted to a probit analysis using the XL-Stat Excel module (Addinsoft, France) for estimating LD values. Resistance ratios (RR_50_) to each insecticide were computed from LD_50_ values obtained for each line as compared to the susceptible strain. Resistance levels of F1 and F2 individuals from each line were obtained following the same procedure.

Cross-resistance profiles of F2 individuals from each line to all insecticides were then evaluated using single-dose bioassays. For each insecticide, the dose was calibrated in order to obtain a mortality ranging from 20% to 40% in the corresponding F2 line. Doses used were as follows: deltamethrin 0.05% for 60 min, bendiocarb 0.5% for 60 min, fenitrothion 0.1% for 45 min. At least 4 batches of 20 three days-old non blood fed females were used per line and insecticide. Mortality was recorded after a 24h recovery time and data were expressed as mean % mortality ± SD. Resistance levels to each insecticide were compared across lines using a generalized linear mixt model (binomial family) using R version 3.5.2 (R Core development Team).

### Deep targeted DNA-sequencing

#### Sample preparation

Deep targeted DNA pool sequencing was used to search for genomic variations associated with insecticide resistance in each line. Genomic DNA was extracted from 2 batches of 50 adult females from each condition (F0 GuyR, F0_LD80_, F2_LD25_ and F2_LD75_, see Fig. 6) using the PureGene kit (Qiagen) following manufacturer’s instructions. DNA extracts obtained from each batch were quality-checked on agarose gel, quantified using the Qubit dsDNA Broad Range kit (Qiagen) and mixed in equal quantity in order to obtain a single genomic DNA extract representative of 100 individuals for each condition.

#### Capture libraries preparation and sequencing

The capture of genomic regions of interest was based on the SureSelect® target enrichment system (Agilent Technologies). Capture library was designed based on Aaeg L3 genome assembly and Aaeg L3.3 annotation and consisted in 54,538 overlapping RNA probes of 120 bp. Among them, 32,494 probes targeted the exons and 1.5 kb upstream regions of 336 candidate genes with a mean coverage of 4X. The remaining 22,044 probes targeted 7,348 unique 220 bp intergenic regions equally spread over *Ae. aegypti* genome. Candidate genes included all known detoxification enzymes (cytochrome P450s, glutathione S-transferases, carboxylesterases, UDP-glycosyltransferases) together with other enzymes potentially involved in insecticide biodegradation pathways and insecticide target proteins. Intergenic regions were defined in order to cover > 95% of *Ae. aegypti* genome using the following criteria: target region size = 220 bp; optimal distance between 2 regions = 150 kb ± 10 kb; region distance to any annotated gene > 5 kb; avoid repeated and redundant regions; avoid regions with GC richness > 70% or single nucleotide richness > 50%; avoid regions with undefined nucleotides (N); do not consider supercontigs < 150 kb; avoid regions located within 75.5 kb of supercontig boundaries. All genomic regions targeted by the study are detailed in S4 Table. Capture was performed with the SureSelect^XT^ Reagent kit (Agilent Technologies) following the ‘SureSelect^XT^ Target Enrichment System for Illumina Paired-end Sequencing Library’ protocol vB.4. Briefly, 3 µg of genomic DNA from each sample were fragmented using a Bioruptor (Diagenode), purified, ligated to adaptors and amplified by PCR using Herculase II DNA polymerase (Agilent Technologies). After QC of library size and quantity, libraries were hybridized to biotinylated baits and purified using Dynal MyOne streptavidin beads (Invitrogene). Captured DNA fragments were amplified, purified and multiplexed before sequencing. Sequencing was performed on an Illumina NextSeq500 and generated more than 300 million 75 bp paired reads with an average of 23.3 million reads per sample. Reads were assigned to each sample (unplexing) and adaptors were removed. Reads quality was checked for each sample using FastQC (http://www.bioinformatics.babraham.ac.uk/projects/fastqc) and reads were loaded into Strand NGS v3.1.1 (Strand Life Science) for further analyses.

#### Reads mapping and filtering

In order to minimize false positives arising from mapping bias in high-redundancy and low complexity regions, CNV were only identified from coding regions. Reads were mapped against all Aaeg L5 exons using a padding of 35 bp and the following parameters: minimum 90% identity, maximum 5% gap, mean insert size of 167 ± 30 bp, mismatch penalty = 4, gap opening penalty = 6, gap extension penalty = 1, clipping penalty = 5, min align read length = 30, ignore reads with more than 5 matches, trim 3’ end if base quality < 25. Reads were then filtered on their quality metrics and mapping quality using the following criteria: mean read quality ≥ 28, N allowed ≤ 2, alignment score ≥ 90, Mapping quality ≥ 40, read length ≥ 35, remove non primary multiply mapped reads, remove inter-chromosomal split reads. Finally, mate missing, translocated and duplicated reads were removed. For polymorphisms analysis, reads were mapped against the whole Aaeg L5 genome in order to consider both genic and intergenic target regions and maximize genome coverage for the detection of selection signatures. The same mapping and filtering parameters as for CNV were applied.

#### CNV detection

The coverage of all exonic regions was computed and only regions showing a mean coverage between 30 and 800 reads/bp in all samples and a length > 45 bp were retained. For all remaining regions, the normalized coverage was used for computing a relative copy number variation in each sample as compared to a common reference obtained from all samples. Relative copy number values were averaged per gene and centering and dimensionality reduction was applied to minimize stochastic variation. For each line, genes were considered affected by CNV associated with insecticide resistance if their CNV profile satisfied the following conditions: CNV(F0_LD80_ - F0)>0.3 AND CNV(F0_LD80_ - F2)>0 AND [CNV(F2_LD25_ - F2)>0.2 OR CNV(F2_LD75_ - F2)>0.2]. Basically, CNV associated with resistance were expected to increase from F0 to F0 survivors, decrease from F0 survivors to F2 after crossing with the susceptible strain, and increase in F2 survivors. No dose response condition was applied for CNV in order to minimize the confounding effect of target site mutations.

#### Polymorphisms and selection signatures

Variants were call against the whole Aaeg L5 genome using the following parameters: coverage > 30 in all conditions, confidence score cut off = 100, ignore loci with homopolymer stretch > 4, ignore loci with average base quality ≤ 15, ignore loci with strand bias ≥ 50 and coverage ≥ 50, ignore reads with mapping quality ≤ 20, ignore variants with less than 4% supporting reads. Among all variants called only those polymorphic among our conditions (*i.e.* showing ≥ 5 % variation in at least two conditions) were retained and their genic effects were computed.

Associations between polymorphisms and resistance to each insecticide were assessed by combining an allele frequency filtering approach with an *F*_ST_-based approach.

The frequency filtering approach was based on the expected resistance allele frequency variations across conditions taking into account their initial frequency. Frequency thresholds used are described in detail in Table 1. Basically, the frequency of alleles positively associated with resistance was expected to increase from unexposed F0 individuals to F0 survivors, decrease from F0 survivors to unexposed F2 individuals (following crossing with the susceptible strain), and increase again in F2 survivors in association with the insecticide dose. Different initial allele frequency thresholds were used for identifying alleles associated with deltamethrin (≥30% in F0 Guy-R) and those associated with bendiocarb and fenitrothion resistance (≥15% in F0 Guy-R) to account for the higher deltamethrin resistance level of the initial composite population. The frequency of deleterious alleles (*i.e.* those negatively associated with resistance) was expected to behave reciprocally.

**Table 1.**
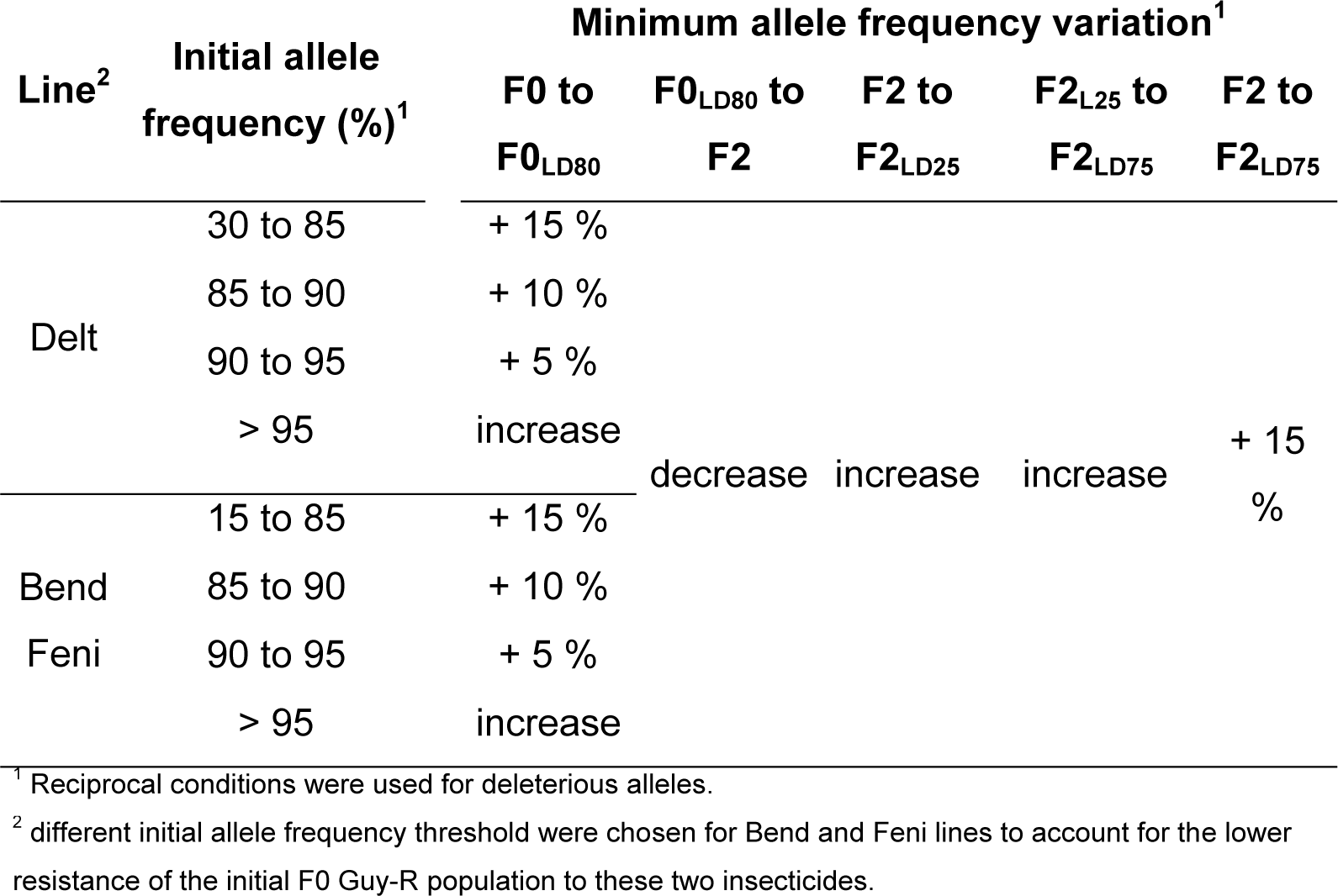
Conditions used for identifying polymorphisms associated with resistance

The *F*_ST_-based approach aimed at assessing departure from neutrality using the Bayesian method implemented in BayeScan version 2.1 [59]. Because substitutions and deletions may have different probability of occurrence, only substitutions were considered for this analysis. For each insecticide line, 2 analyses were run separately. The first one contrasted allele frequencies in F0 samples (unexposed and insecticide survivors: F0 Guy-R and F0_LD80_) and the second one in F2 samples (unexposed and survivors to each insecticide dose: F2, F2_LD25_, F2_LD75_). The Markov chain Monte Carlo (MCMC) algorithm was run with prior odds of 1. 10. The proposal distributions for parameters were adjusted by running 20 short pilot runs of 2,000 iterations. A burn-in period of 100,000 iterations was used and the posterior probabilities were estimated from the following 500,000 iterations (10,000 iterations samples every 50). Genomic regions showing low Bayscan *Q*-values in both F0 and F2 comparisons and including differential polymorphisms identified from the frequency filtering approach were considered as under selection in association with insecticide resistance.

### Kdr mutations genotyping

Allelic frequencies for the three kdr mutations (V410L, V1016I and F1534C) initially present in the F0 Guy-R composite population were obtained from F0 and F2 samples of each line based on reads data. In order to validate allele frequencies obtained from targeted pool DNA-seq, the two kdr mutations V1016I and F1534C were genotyped on individual mosquitoes from the initial F0 Guy-R population (F0) together with F0 and F2 samples of the Delt line. Total genomic DNA was extracted using cetyl trimethyl ammonium bromide chloroform/isoamyl alcohol from 30 non blood fed females per condition as described in [60]. Individual genotypes for each kdr mutations were obtained by qPCR high resolution melt curve analysis using 0.15 ng of genomic DNA per reaction as described in [20].

### Data availability statement

The sequence data from this study have been deposited to the European Nucleotide Archive (ENA; http://www.ebi.ac.uk/ena) under the accession number PRJEB30945.

## Acknowledgments

We thank Dr. Tristan Cumer, Dr. Frederic Boyer, Prof. Laurence Despres for critical comments on the manuscript.

## Author contributions

Conceived and designed the experiment: JPD. Performed experiments: FF, BL, MM, LG, TG, FL, SR. Analyzed the data: JC, FF, BL, SS, SR, JPD. Collected biological material: FF, ID. Wrote the manuscript: JC, JPD.

## Supplementary materials

**S1 Fig. Cross-validation of kdr mutation frequencies obtained by mass sequencing and individual genotyping.** Blue dots show the allele frequency obtained by targeted DNA-seq with the total read coverage indicated for each condition. Empty blue dots designate conditions for which the total read coverage was < 30. Triangles designate allele frequencies obtained by qPCR genotyping with the number of genotyped individual indicated for each condition.

**S1 Table. CNV data set.** This table shows the mean relative copy number of each gene across all F0 and F2 conditions. Relative gene copy numbers were obtained by comparing the normalized coverage of each target region to a common reference made from all samples. Only regions showing a mean coverage between 30 and 850 in all samples were considered. Relative copy numbers were then averaged per gene and centered-reduced in order to minimize stochastic variations. Genes affected by CNV associated with resistance to each insecticide are indicated.

**S2 Table. Polymorphism data set overview.** This table describes the overall polymorphisms data set. Polymorphisms counts and percentages are given for the following categories: passing QC filters and polymorphic across samples, differential in each line, non-synonymous differential in each line.

**S3 Table. Polymorphism data set.** This table describes all variations passing QC filters and polymorphic across samples. For each variation, the following attributes are listed: location (chromosome, start, end), reference and variant allele, polymorphisms type, target region type, variant allele frequency in each condition, differential based on allele frequency filtering (Yes or No for each line), Bayescan -Log_10_ Q value (for F0 and F2 condition in each line), genic effect based on AaegL5.1 annotation (effect, affected gene accession and description, AA change, cDNA position/effect, protein position/effect).

**S4 Table. Genomic regions targeted by DNA-seq.** This table describes all genomic regions targeted by DNA-seq. For each target region, the following information are shown: AaegL5 location (chromosome, start, end, length), AAegL3 location (supercontig: start-end, length), marker type (genomic “marker”, gene “promoter” or “gene”), short description, protein family.

